# Surface area calculations of lamellar support respiratory function of trilobite exopodites

**DOI:** 10.64898/2026.02.02.702636

**Authors:** Sarah R. Losso, Federica Vallefuoco, Igino Foglia, Léo Laborieux, Ana Belén Muñoz-García, Javier Ortega-Hernández

## Abstract

Trilobites had biramous appendages with an inner endopodite (walking leg) and outer exopodite (gill) connected to the body through the protopodite (limb base). Whereas both endopodite and protopodite were involved in both locomotion and feeding, the exopodite has been subject to various functional interpretations including respiration, ventilation and swimming. Evidence from sites with exceptional fossil preservation indicate that trilobite exopodites show substantial variability in terms of the number and size of their articles, lamellae and setae, but the implications of this morphological diversity have never been investigated. Here, we created anatomically correct 3D models of exopodites in *O. serratus* and *T. eatoni* to calculate the SA of the lamellae and explore its relationship with body size. Our results indicate a large SA for *O. serratus* at 16,589 mm^2^ compared to the 2,159 mm^2^ for the much smaller *T. eatoni*. We also calculated lamellar SA for nine additional trilobite species with exceptionally well-preserved appendages based on lamellar measurements. The results indicate that lamellae SA of trilobites increased exponentially with overall body size. Trilobite data follows the same trendline of gill SA/biomass observed in extant species and thus supports the interpretation of their exopodites as respiratory structures despite substantial variation in morphology.

## Introduction

Aerobic respiration is one of the most fundamental functions of metazoans, required to exchange oxygen and produce metabolic energy. The respiratory function of aquatic euarthropods is often performed by gills which serve as the location of diffusion of oxygen into the body to be distributed to cells. Due to their varied lifestyles and diverse requirements, euarthropods have evolved a broad variety of gill morphologies and sizes that follow the enormous ecological versatility of these organisms (Hughes et al., 1969; Suzuki and Bergström, 2008; Suzuki et al., 2008; Glazier and Paul, 2017; Liu et al., 2020, Liu et al., 2023; Fig. 1). While many modern aquatic euarthropods often have limbs specialized for respiration or locomotion (Fig. 1g – i), most Early Paleozoic representatives had biramous appendages with an endopodite (e.g. inner branch, or walking leg) and a lamellae bearing exopodite (i.e. outer ramus) (Boxshall, 2004) (Fig. 1a-f). However, the precise function of the exopodites among Paleozoic euarthropods has been controversial. The main functional hypotheses put forward include their involvement in swimming based on their flattened morphology (Haug and Haug, 2016; Zeng et al., 2017), as ventilatory structures based on estimates of overall surface area (SA) compared with extant euarthropods (Bergström, 1969; Suzuki and Bergström, 2008; Suzuki et al., 2008), or respiration and gas exchange based on the presence of dumbbell-shape lamellae akin to the gills of living species (Hou et al., 2021).

**Figure 1.**
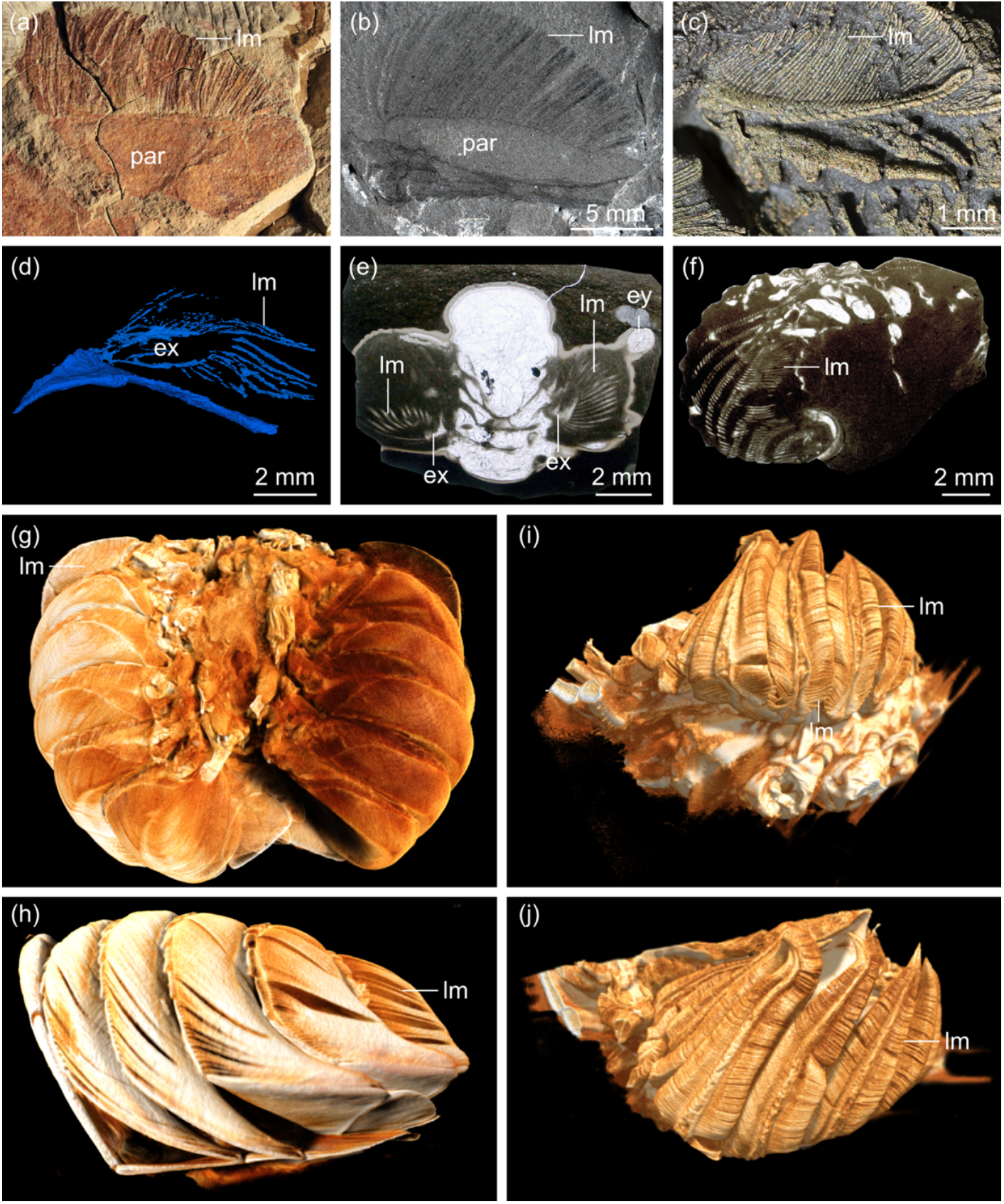
Exopodite and gill morphology in aquatic euarthropods. **(*a*)** Exopodite of *Redlichia rex* from Emu Bay (SAM P5486; Cambrian, Stage 2; Australia; Holmes et al., 2020). **(*b*)** Exopodite of *Olenoides serratus* from the Burgess Shale (USNM PAL 188574; Cambrian, Wuliuan; Canada; Losso et al., 2025). ***c*)** Exopodite of *Triarthrus eatoni* from Beecher’s Trilobite Bed (GLAHM 163103; Ordovician, Katian; USA; Hou et al., 2023). **(*d*)** Exopodites of *Dalmanites* sp. from the Herefordshire biota (OUMNH C.29611; Silurian, Homerian; USA; Siveter et al., 2021). **(*e*)** Exopodites of *Ceraurus pleurexanthemus* from the Walcott-Rust Quarry (MCZ:IP:104973; Ordovician, Katian; USA; Losso and Ortega-Hernández, 2024). **(*f*)** Exopodites of *Flexicalymene* sp. from the Walcott-Rust Quarry (USNM PAL 68379; Ordovician, Katian; USA; Losso and Ortega-Hernández, 2024). **(*g – h*)** Micro-CT scanned dissected gills from modern *Limulus polyphemus*. **(*g*)** Dorsal view. **(*h*)** Lateral view. **(*i – j*)** Micro-CT scanned dissected gills from modern *Cancer borealis*. **(*g*)** Lateral view. **(*h*)** Dorsal view. Abbreviations: ey, eye; lm, lamella; par, proximal article.

SA estimates have previously only been calculated for two trilobites, the mid-Cambrian (Wuliuan) *Olenoides serratus* at 153mm^2^ for one exopodite (Suzuki et al., 2008), and the Late Ordovician (Katian) *Triarthrus eatoni* at 2,067mm^2^ for the entire body (Hou et al., 2023). However, trilobite exopodites have a broad range of morphological variation in terms of the number and size of articles, lamellae, size of limbs relative to the body, and number of appendage pairs (Ortega-Hernández et al., 2013; Losso et al., 2026). The SA of gills is an important control on the amount of oxygen that can diffuse into the body (Gray, 1957; Hughes et al., 1969; Glazier and Paul, 2017), and thus directly affect the amount of energy available to the individual.

Here we analyze the SA of exopodites in 11 trilobite species spanning a broad range of biomass. We demonstrate that exopodite surface area increased exponentially with biomass owing to an elongation of the lamellae in larger taxa. Our results indicate that trilobites had exopodite SA to biomass ratios comparable to those of modern aquatic crustaceans and the Atlantic horseshoe crab. Our fossil data supports the interpretation of trilobite exopodites as having a respiratory function as they show the same relationship between gill SA and biomass as that observed in modern aquatic euarthropods.

## Methods

### Specimens and micro-CT scanning

Studied fossil specimens are housed in the Smithsonian Institution (Washington D.C., USA; USNM), The Hunterian Museum, University of Glasgow (Glasgow, United Kingdom; GLAHM), South Australian Museum (Adelaide, Australia; SAM), Museum of Comparative Zoology (Cambridge, Massachusetts, USA; MCZ), and Oxford University Museum of Natural History (Oxford, United Kingdom; OUMNH). One specimen of the Atlantic horseshoe crab, *Limulus polyphemus* Linnaeus, 1758, was purchased from Woods Hole Oceanographic Institute with the gills dissected from the opisthosoma and fixed in 10% Neutral Buffered Formaldehyde, then stained in 1.25% iodine for four days in preparation for micro-Computer Tomography (micro-CT) scanning. One specimen of the Jonah Crab, *Cancer borealis*, was purchased from New Deal Fish Market (Cambridge, Massachusetts, USA) and euthanized through spiking. The gills were dissected from the body, stained in 1.25% iodine in water for 15 hours. The gills of both *L. polyphemus* and *C. borealis* were then micro-CT scanned using a Bruker Skyscan 1273 at the Harvard Digital Imaging Facility. The *L. polyphemus* specimen was scanned using a voltage of 80kv, wattage of 300uA, and a 0.5mm aluminum filter. The *C. borealis* specimen was scanned using a voltage of 115kv, wattage of 69uA, and a 0.1mm aluminum plus 0.038mm copper filter. The scans were reconstructed as TIFF stacks in NRecon (Bruker Corporation) and visualized in Dragonfly 19 4.0 (Object Research Systems, Montreal, Canada).

### Three-dimensional modeling

Lamellar SA for two well-preserved species (*Olenoides serratus* and *Triarthrus eatoni*) was calculated by three-dimensional modeling (Tier 1 path, Supplemental Fig. 1) to get highly accurate results and compare with other methods (see below). Three-dimensional models of exopodites were created in Shapr3D for two taxa based on the best-preserved limbs available for study, namely *Olenoides serratus*, USNM 188574 (Fig. 1b; Supplemental Fig. 1; Losso et al., 2025) and *Triarthrus eatoni*, GLAHMN 163103 9^th^ thoracic appendage (Fig. 1c; Hou et al., 2023). Lamellae dimensions were measured on USNM PAL 188574 showing the appendage in dorsal view and a unique specimen, USNM PAL 65514, showing the cross section of the lamellae. The thickness of the exopodite articles is not directly measurable in any specimens of *O. serratus*, but was modeled as being equivalent to the height of the lamellae as seen in other trilobites (Hou et al., 2021; Losso and Ortega-Hernández, 2024). The presence of dumbbell-shaped lamellae, their dimensions and number are however clearly visible in multiple specimens (see Losso et al., 2025). The detailed morphology of the lamellae of *T. eatoni* have been extensively studied recently (Hou et al., 2021, Hou et al., 2023), and the exopodite was modeled as thin articles with a circular cross section based on the available material.

The 3D models were imported into Ansys (Ansys Academic, Release 2020 R2) and exopodite length was scaled to match the measured size in specimens (Fig. 2a-c). All the lamellae were selected, and the SA was calculated. The SA was then doubled to find the total for each limb pair. To find the total lamellar SA along the body for each taxon, the decrease in size of limb pairs were measured or estimated, then used to scale the modeled SA and summed (Supplemental Table 1). The isolated appendage of USNM 188574 is similar in size to the anterior thoracic exopodites of USNM 65521, a specimen that is 67.8 mm long.

**Figure 2.**
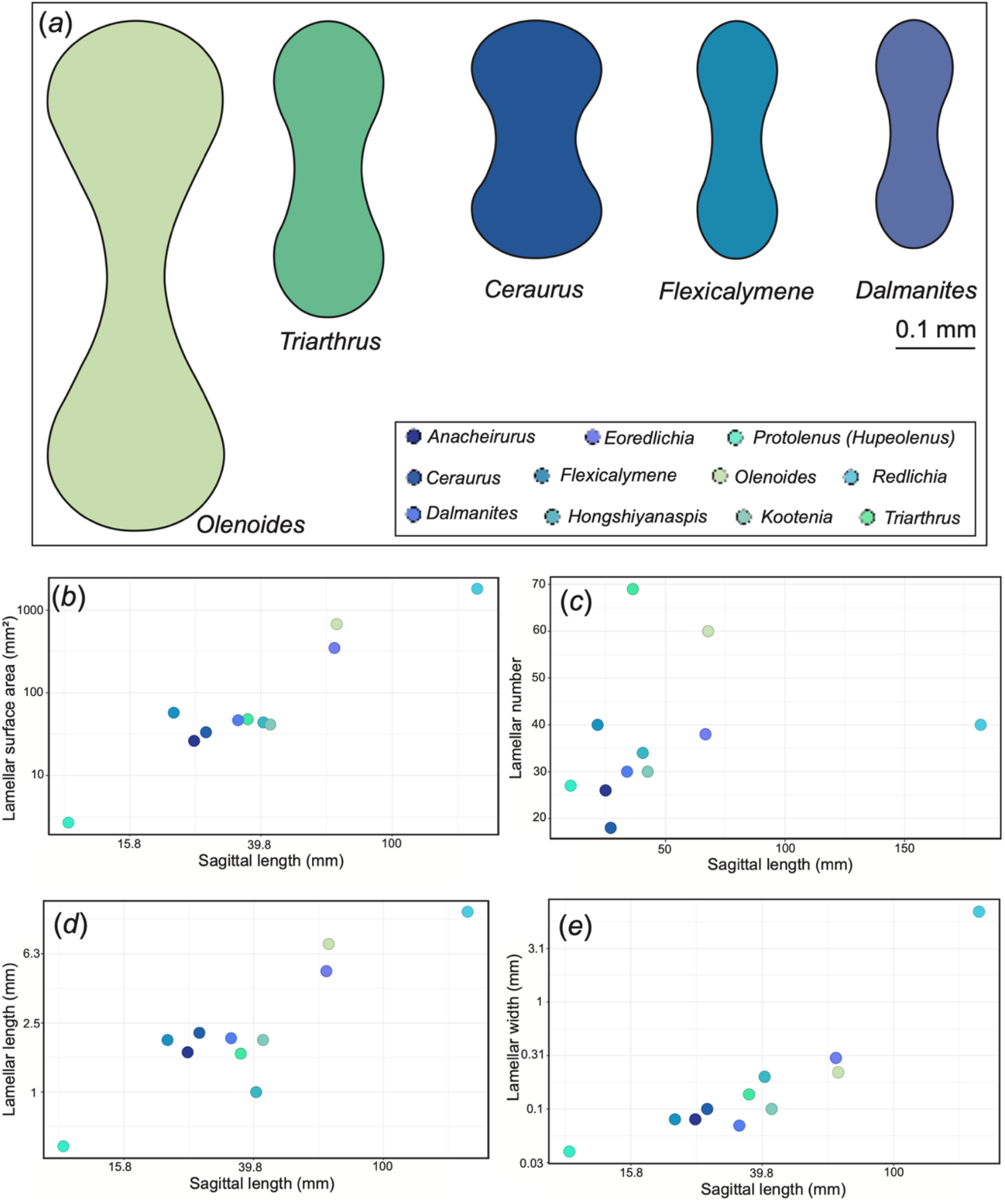
Measurements of trilobite lamellae. **(*a*)** Diagrams of lamellae in five trilobites with well-preserved exopodites showing dumbbell shaped cross section associated with gas exchange. **(*b*)** Log-log plot of lamellar surface area compared to exoskeleton sagittal length. **(*c*)** Lamellar number compared to exoskeleton sagittal length. **(*c*)** Lamellar number compared to exoskeleton sagittal length. **(*d*)** Log-log plot of length of one lamella compared to exoskeleton sagittal length. **(*e*)** Log-log plot of width of one lamella compared to exoskeleton sagittal length.

A single lamella of *Olenoides serratus* was modeled with the dumbbell shaped cross section and compared with an oval cross section with the same right and maximum width to investigate the impact on surface area. The dumbbell shape increased surface area by 2.3% (Supplemental Table 1).

### Surface area calculations through lamellar measurement

We measured the lamellar dimensions of three taxa with high quality preservation in cross section, *Ceraurus pleurexanthemus, Flexicalymene* sp., and *Dalmanites* sp. (Figs. 1d-f, 2a; Supplemental Table 1; Tier 1 path, Supplemental, Fig. 1; Supplemental Fig. 2). The SA of each exopodite was calculated by measuring the perimeter length of the lamellar cross section multiplied by the lamellar length, then by the total number of lamellae per exopodite. The resulting lamellar SA for one exopodite was multiplied by two to account for a complete limb pair. The SA for each appendage pair was then calculated for the entire body by measuring the axial tergite length and scaling the limb pair SA along the body. *Flexicalymene* sp. and *C. pleurexanthemus* appendages are only known from thin sections, so exoskeletal features (e.g. glabella and tergite length) were measured to compare with prone specimens.

Trilobite lamellae are typically preserved in dorsal view (e.g. *Redlichia rex* in Fig. 1A), and thus information about their cross section is not readily available. Consequently, we could only directly measure lamellar length and width for the remaining six species. Lamellar height ranges between 13.6 – 15.4% length for *C. pleurexanthemus, Flexicalymene* sp., and *Dalmanites* sp. (Supplemental Table 1). For the six species where lamellar height was not measurable, it was estimated as 15% of the length base on proportions seen in the other better-preserved taxa. The perimeter length of oval cylinder was calculated, then increase by 2% to account for the dumbbell shape (Supplemental Table 1). The resulting SA was multiplied by lamellae length, then doubled to account for a full limb pair. Total SA for the body was calculated as above (Supplemental Table 1).

### Biomass calculation

Biomass was estimated by finding the length and width of each specimen to create an oval. The width of the axial lobe was used to estimate thickness based on the anatomical data reported in Losso and Ortega-Hernández (2024) and used to calculate the volume for most taxa. As *Flexicalymene* has a more convex exoskeleton, the thickness was estimated as two third the total width to account for the greater depth. A density of 1.1g/cm^3^ was used to estimate the biomass for each taxon (Hou et al., 2023). SA/biomass was calculated using the total gill SA (mm^2^) and biomass (g) to compare with measurements in extant animals (Supplemental Table 1).

## Results

### Trilobite lamellae compared to biomass

Trilobite lamellae show a dumbbell shaped cross section in all instances where it is preserved (Fig. 2a). They exhibit variation in lamellar height and width, with *Olenoides serratus* being the tallest and *Ceraurus pleurexanthemus* being the widest proportionally (Fig. 2a). Number of lamellae ranges from 18 (*C. pleurexanthemus*) to 69 (*Triarthrus eatoni*) and does not correlate with specimen size (Fig. 2c). Lamellar length ranges from 0.487 mm in *Protolenus (Hupeolenus)* sp. to 11.02 mm in *Redlichia rex* (Fig. 2d).

Our results indicate that trilobites follow an exponential increase in lamellar SA relative to exoskeletal sagittal length, but interestingly the number of lamellae and limbs is not directly correlated with sagittal length (Fig. 2b, c; Supplemental Table 1). Lamellae length increases linearly with larger body sizes (Fig. 2d), whereas lamellar width increases much more slowly (Fig. 2e). For example, *Redlichia rex* is the largest trilobite with preserved appendages and its exopodites bear 40 lamellae, but these are drastically longer and wider than those found in smaller species (Fig. 2). *Olenoides serratus* has a high SA per exopodite, in part from having 60 lamellae. Although *Triarthrus eatoni* has up to 69 lamellae, it has similar SA to the similarly sized taxa *Eoredlichia intermediata, Dalmanites* sp., and *Hongshiyanaspis yiliangensis* (Fig. 2b, c).

### Lamellae surface area compared to biomass

Modern marine euarthropods exhibit a broad range of gill SA, from 1,435.2 to 294,172 mm^2^ depending on clade and size (Supplemental Table 2). Within the included species, marine brachyurans have SA(mm^2^)/biomass(g) ranging from 500 – 1367, whereas thalassinids can have a lower ratio between 256 to 1,043 (Fig. 3a; Supplemental Table 2). Xiphosura, represented only by the Atlantic Horseshoe Crab, has a similar ratio as brachyurans at 1,124.38mm^2^/g (Fig. 3a; Supplemental Table 2). Trilobites show a lower but overlapping ratio from 174.62 – 759.48mm^2^/g (Fig. 3a).

**Figure 3.**
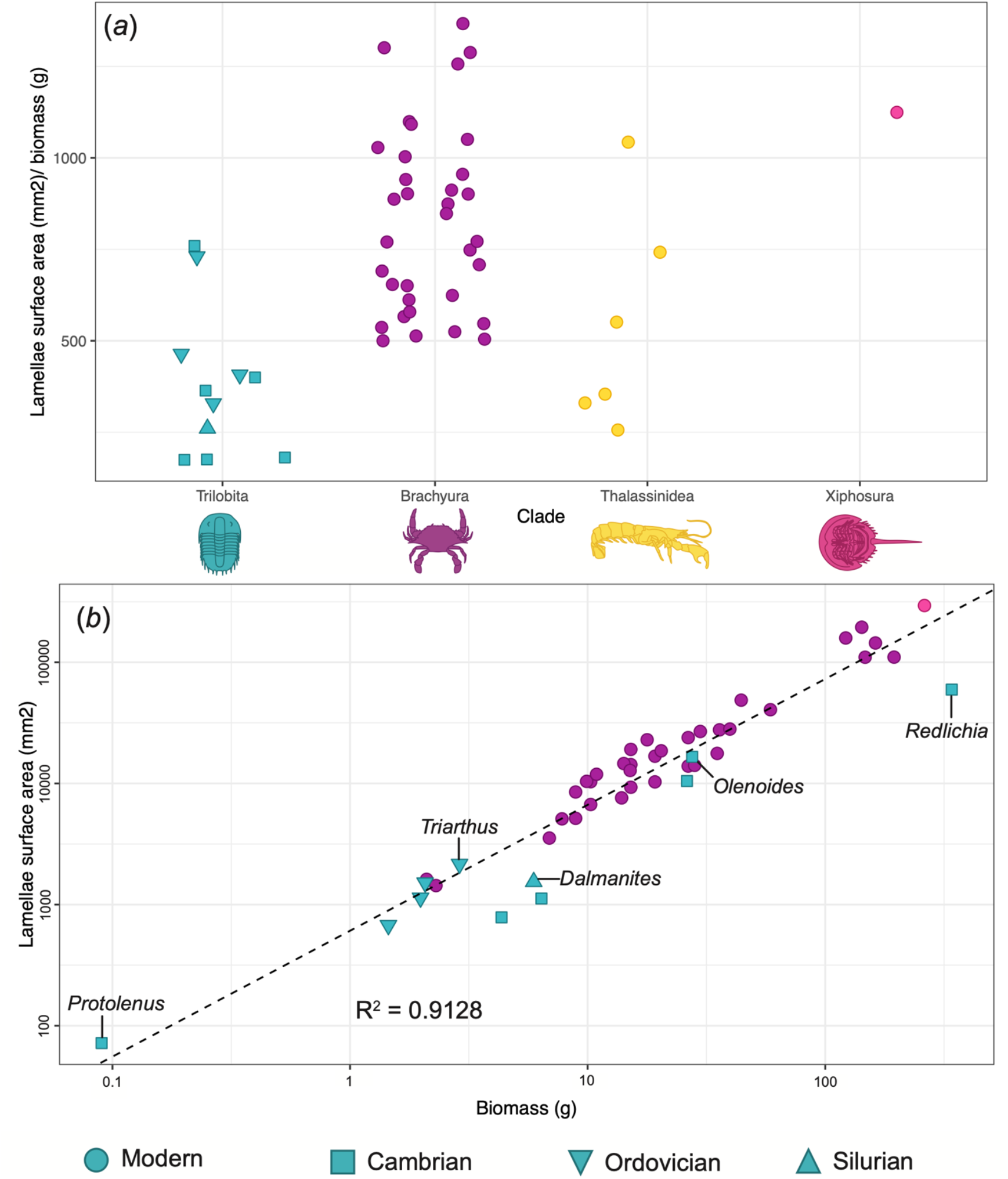
Gill surface area to biomass in aquatic euarthropods. **A**, Gill surface area/biomass by clade (Supplemental Table 2). **B**, Log-log plot of bivariate plot of surface area to biomass color coded by clade. Thalassinidea data is only reported as a ratio and cannot be included in plot B.

The variation in modern gill SA is strongly associated with biomass and increases exponentially (Fig. 3b). At smaller sizes, the increase in SA is strongly correlated with biomass, but over 122g, the pattern becomes more complex (Supplemental Fig. 2b). Species included in the analyses were weighed using various methods, but the variation in gill size compared with biomass is not explained by this (Supplemental Fig. 2a). *Libinia emerginata, Libinia dubia*, and *Menippe mercenaria* are large crabs, but with comparatively low gill SA (Fig. 3b). The largest modern euarthropod included in the analysis was *Limulus polyphemus* at 261.63g (dry weight), with a SA/biomass ration that follows the trend of smaller species (Fig. 3b). All trilobites except *Redlichia rex* follow a similar trend as the modern taxa, with some falling slightly lower. The most notable deviation is *R. rex*, the largest trilobite included in the analysis, with a substantially lower surface area and an estimated biomass larger than *L. polyphemus*.

## Discussion

### Relationships between exopodite size and morphology

Although trilobite species exibit lamellae with a dumbbell shaped cross section, there is variation in their width and height. The Ordovician *Ceraurus pleurexanthemus* has wider lamellae compared to *Flexicalymene* and *Dalmanites*, which both have a similar height (Fig. 2a). The rapid increase in lamellar SA with size is accomplished through elongation of the filaments (Fig. 2d) rather than increase in the number of lamellae (Fig. 2c) or width (Fig. 2e). Thalassinidean shrimp also increase gill surface area through changes in filament shape rather than increased numbers (Astall et al., 1997). Increased length would result in faster growth of SA rather than increased width. *Ceraurus* has a lamellar SA of 1.8568mm^2^ vs. *Flexicalymene* at 1.44mm^2^ and is 1.6x as wide as the latter (Supplemental Table 1). A 1.6x increase in length of the lamellar in *Flexicalymene* would result in an SA of 2.53mm^2^. The distance between each lamella in trilobites is usually roughly equal to the maximum width (Hou et al., 2023; El Albani et al., 2024; Losso and Ortega-Hernández, 2024), with a notable exception in *Dalmanites* (Siveter et al., 2021). As water flows between the lamellae, increased width may decrease the spacing or require a longer exopodite to allow for equal spacing to be maintained. In the former instance, this would impact the pattern and speed of water flow between lamellae (see Hou et al., 2023), and the latter could expose the more delicate exopodites from underneath the exoskeleton as seen in *Triarthrus eatoni*. Variation in SA/biomass within a clade may reflect respiratory needs depending on environment (Gray, 1957; Johnson and Rees, 1988; Watson-Zink, 2021), burrowing resulting in exposure to anoxia (Astall et al., 1997), and morphological pressure for streamlined body plans (Wootton et al., 2015). The exopodite morphology of *Triarthrus eatoni* has been suggested to maximize lamellar SA for oxygen exchange, but the low oxygen uptake compared to modern fish and crustaceans may have requires a less energetic lifestyle (Hou et al., 2023). Our work confirms the maximization of lamellar SA in *T. eatoni* (Fig. 3b) compared to other trilobites which make have lived in more oxygenated water.

*Redlichia rex* displays low lamellar SA compared to other trilobites and modern marine euarthropods for its size (Fig. 3b), which could result from several factors. No specimen of *R. rex* is known with attached limbs, the disarticulated limbs are associated with the taxon because of the large size compared to any other trilobites from the Emu Bay Shale (Holmes et al., 2020). The limbs were measured and compared to an articulated specimen, but may not reflect the true size relationship. Alternatively, if this is an accurate representation of the limb size, *R. rex* may have a lower metabolic requirement depending on lifestyle (Hou et al., 2023) or gain additional oxygen through the ventral side of pleural lobes (Suzuki and Bergström, 2008). In some fish species such as *Clupea harengus*, there is a steep slope between gill surface area and weight, but the gill areas increases more slowly at large sizes (Pauly and Müller, 2025). A similar trend may be found in euarthropod gills, but more data of large individuals is needed to evaluate this.

### Trilobite exopodites as gills

The lamellar SA of exopodites has direct implications for whether these structures were used for respiratory or ventilatory functions in trilobites (Suzuki and Bergström, 2008; Suzuki et al., 2008). The similarity between our estimates for total SA and that of Hou et al. (2023) despite different methods of scaling limbs along the body, supports this as a robust estimate. But the disparity between our estimate of lamellar SA for one exopodite of *O. serratus* compared to that of Suzuki et al. (2008) would have a significant impact on the total SA, resulting in a particularly low ratio of area to biomass. This feature was used by Suzuki et al. (2008) to support a non-respiratory interpretation of exopodites, as they would not allow for enough oxygen exchange given the size of the animal. This is most apparent when comparing to *Limulus polyphemus* which has 294,172mm^2^ for a 261.63g specimen. As noted by (Hou et al., 2023), the SA/biomass ratio of *T. eatoni* between 300 – 450 mm^2^/g -or 407.07 by our calculations-, is within the range of modern decapods, albeit was closer to above tide taxa. Even with the restriction to aquatic euarthropods, SA/biomass of all measured trilobites is comparable to modern taxa (Fig. 3). The ventral surface of the pleural lobes in trilobites may have also acted as a site of oxygen exchange (Suzuki and Bergström, 2008; Suzuki et al., 2008) providing additional surface area. The mud shrimp *Calocaris macandreae* has a low SA/biomass at 256mm^2^/g while *Upogebia pusilla* and *Upogebia stellata* have ratios of 354mm^2^/g and 330mm^2^/g respectively (Astall et al., 1997). As SA scales exponentially with biomass, it is inappropriate to restrict comparisons to the SA/biomass ratio, which will be substantially higher in larger species. Most trilobites are smaller than the brachyurans, but surface area of exopodite lamellae follows the same trends as the modern taxa, supporting the interpretation as gills.

## Acknowledgements

This work was supported by a Research Grant from the Human Frontier Science Program (Ref.-No: RGY0056/2022). We thank Pauline Affatato (University of Columbia, New York, USA) for micro-CT scanning the specimen of *Limulus polyphemus*; Javier Luque (Cambridge University, Cambridge, United Kingdom) for discussing crab morphology imaging; Jared C. Richards (Harvard University, Cambridge, MA, USA) for discussion of data visualization; Walker C. Weyland for help rendering 3D models; James Holmes (Uppsala University and University of New England) and Derek Siveter (University of Oxford, Oxford, United Kingdom) for providing an image of *specimens*; Jessica Cundiff (Museum of Comparative Zoology, Harvard University, Cambridge, MA, USA), Gene Hunt, Conrad C. Labandeira, Doug Erwin, Mark Florence (Smithsonian Institution, Washington D.C., USA) and Neil D. L. Clark (Hunterian Museum & Art Gallery, University of Glasgow, Glasgow, United Kingdom) for facilitating access to specimens.

## Supplemental Information

Table 1. Trilobite exopodite measurements and biomass calculations.

Table 2. Arthropod gill surface area

**Supplemental Figure 1.**
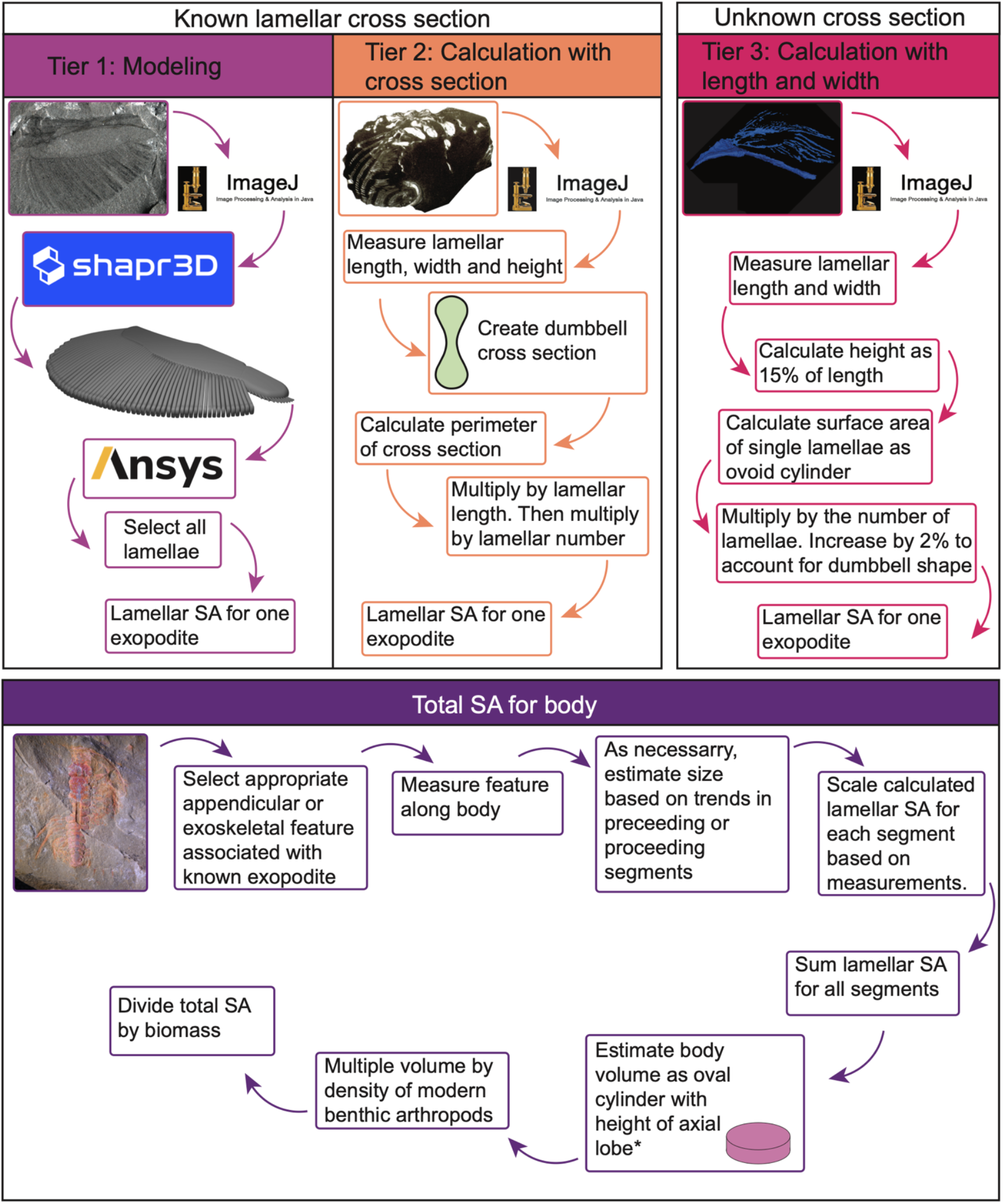
Lamellar surface area calculation pathways.

**Supplemental Figure 2.**
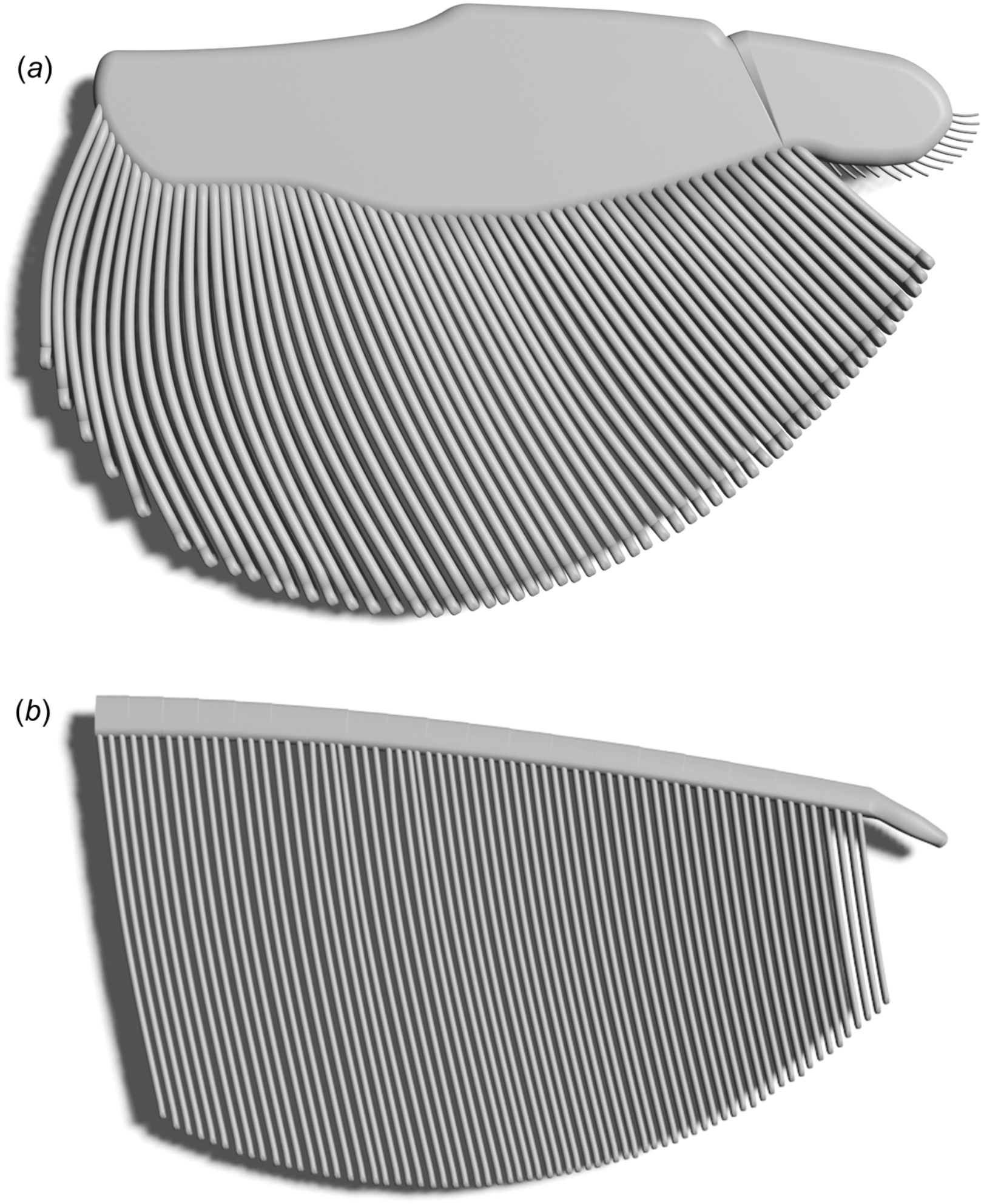
Three-dimensional models of exopodites. A, *Olenoides serratus*. b, *Triarthrus eatoni*.

**Supplemental Figure 3.**
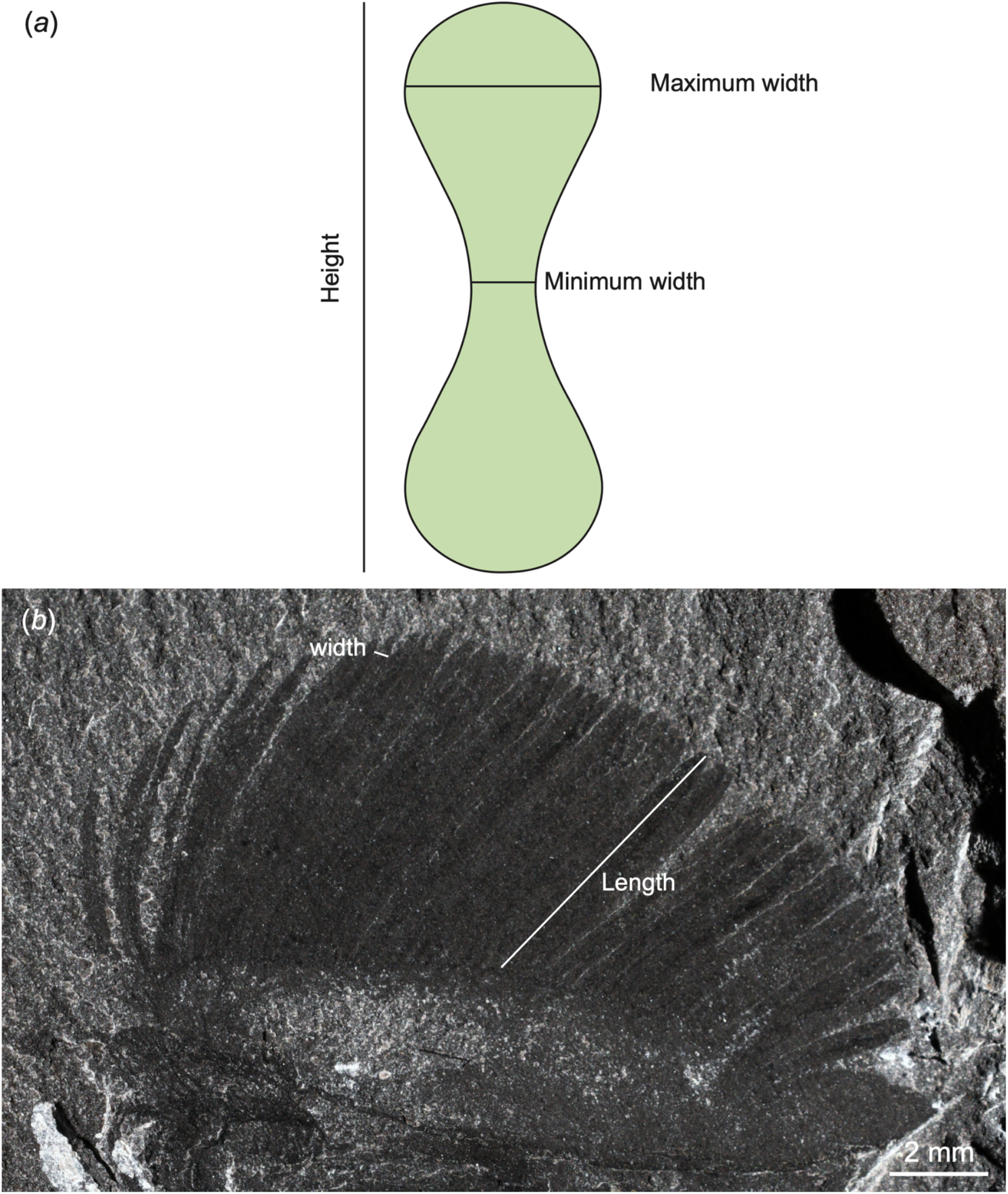
Lamellar terminology and dimensions. A, Lamellar cross section. B, Compressed fossil.

**Supplemental Figure 4.**
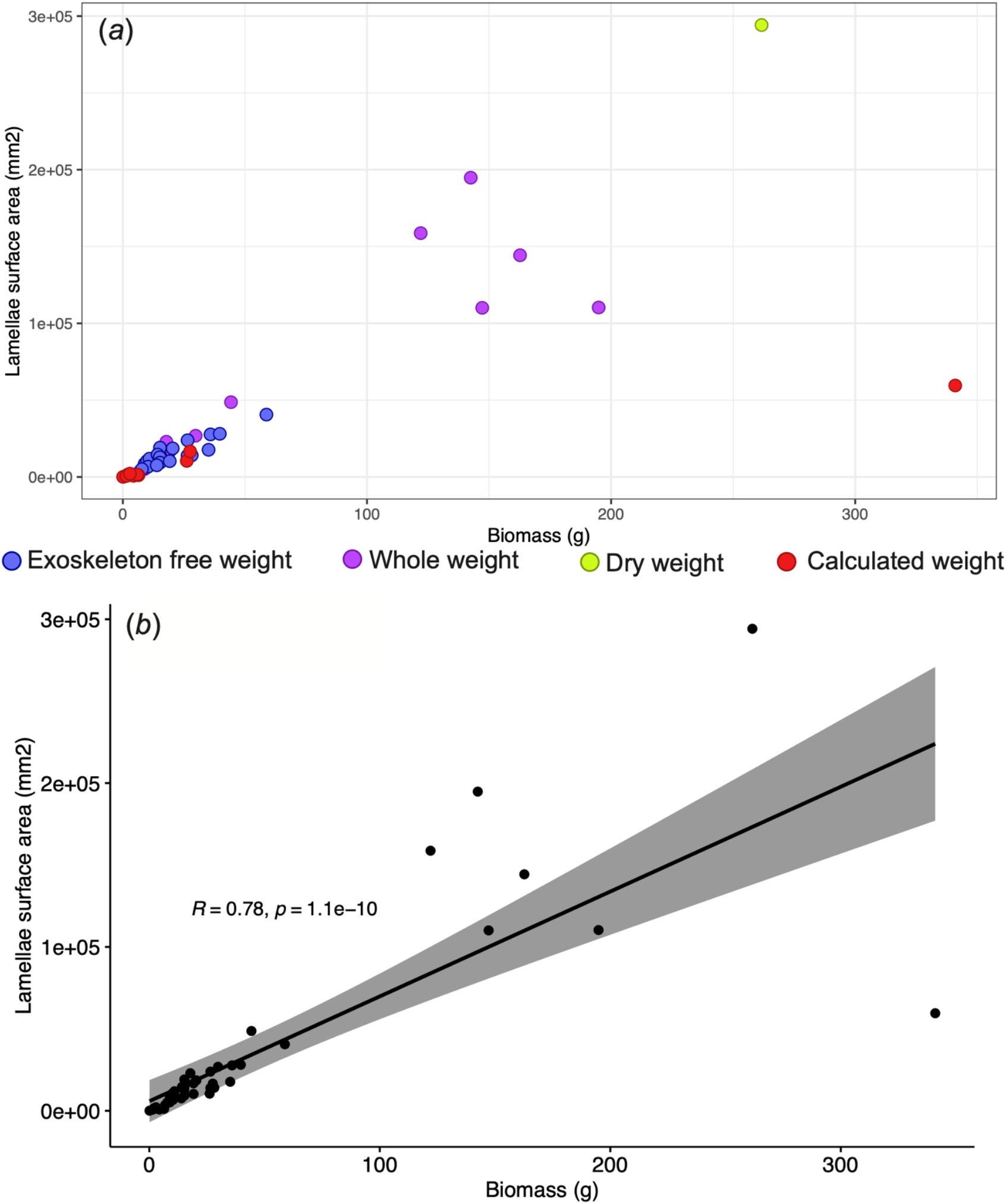
Gill surface area to biomass. A, Weighing method and Pearson’s correlation of lamellar surface area and biomass. B, bivarate plot with 95% confidence interval and linear trend line.

## Notes

### Competing Interest Statement

The authors have declared no competing interest.

